# Protease shaving of *Mycobacterium tuberculosis* facilitates vaccine antigen discovery and delivery of novel cargoes to the Mtb surface

**DOI:** 10.1101/2024.07.02.601718

**Authors:** Bianca A. Lepe, Christine R. Zheng, Owen K. Leddy, Benjamin L. Allsup, Sydney L. Solomon, Bryan D. Bryson

## Abstract

Tuberculosis (TB), caused by *Mycobacterium tuberculosis* (Mtb), is the leading cause of infectious disease death and lacks a vaccine capable of protecting adults from pulmonary TB. Studies have shown that Mtb uses a variety of mechanisms to evade host immunity. Secreted Mtb proteins such as Type VII secretion system substrates have been characterized for their ability to modulate anti-Mtb immunity; however, studies of other pathogens such as *Salmonella* Typhi and *Staphylococcus aureus* have revealed that outer membrane proteins can also interact with the innate and adaptive immune system. The Mtb outer membrane proteome has received relatively less attention due to limited techniques available to interrogate this compartment. We filled this gap by deploying protease shaving and quantitative mass spectrometry to identify Mtb outer membrane proteins which serve as nodes in the Mtb-host interaction network. These analyses revealed several novel Mtb proteins on the Mtb surface largely derived from the PE/PPE class of Mtb proteins, including PPE18, a component of a leading Mtb vaccine candidate. We next exploited the localization of PPE18 to decorate the Mtb surface with heterologous proteins and deliver these surface-engineered Mtb to the phagosome. Together, these studies reveal potential novel targets for new Mtb vaccines as well as facilitate new approaches to study difficult to study cellular compartments during infection.

## Introduction

Tuberculosis (TB), caused by the causative agent, *Mycobacterium tuberculosis* (Mtb), is the leading cause of death due to a single infectious agent^1^. No protective vaccine has been developed that meets the target product profile needed in humans to alter the course of the TB pandemic. Gaps in our understanding of the host-Mtb interactions that facilitate interactions between the host immune system and the bacterium limit the development of more effective TB vaccines, host-directed therapies, and biotechnologies suited to generate new insights about Mtb infection.

Interactions between Mtb proteins and the host underlie both innate and adaptive immune responses. For example, secreted Mtb proteins play a critical role in immune responses. The ESX-1 and 3 secretion systems have been shown to secrete proteins that interact with the endolysosomal system in phagocytes modulating both membrane damage and repair pathways ^2–7^. In addition to modulating innate immune processes, secreted Mtb proteins comprise a significant fraction of the antigenic targets of Mtb-specific T cells that facilitate immune protection during infection^8,9^. These data emphasize the critical role that Mtb proteins, defined by their physical localization in the bacterium, influence immune responses. Technically, the identification of secreted Mtb proteins has been greatly facilitated by optimized biochemical workflows that enable the physical separation of the Mtb secretome from other proteomic fractions ^10–13^ .

Recent studies also emphasize that Mtb proteins anchored in the mycobacterial outer membrane may serve a similar role in bacterial pathogenesis and interactions with adaptive immunity. PPE51, an Mtb outer membrane protein, was recently demonstrated to be anchored in the mycobacterial outer membrane and to facilitate nutrient acquisition, a critical function required for bacterial survival^14^. We and others have also demonstrated that beyond its nutrient acquisition function, PPE51 serves as an antigenic target for CD8+ T cells ^15,16^. Similar observations have been made for other pathogens. In gram-negative bacteria there is a subset of highly conserved outer membrane proteins called porins which play a role in small molecule transport across the membrane and bacterial homeostasis. OmpC and OmpF in *Salmonella enterica* serovar Typhi both induce a robust cell-mediated immune response in mice and humans^17^.

Despite these examples emphasizing a role for outer membrane proteins in Mtb pathogenesis, considerably less is known about the composition of this physical compartment compared to the Mtb secretome^18^. Physical fractionation of the different Mtb compartments has been performed; however, these fractionation methods can be both laborious and equipment intensive due to the requirement of ultracentrifugation, which is difficult to perform in a BSL3 environment. Additionally, ultracentrifugation methods alone are insufficient to define the topology of proteins in the mycobacterial outer membrane. Addressing these gaps in our understanding of the outer membrane proteome would facilitate an improved understanding of Mtb biology as well as potentially facilitate new approaches to characterize the host-pathogen interface. For instance, Olson and colleagues previously demonstrated in *Chlamydia trachomatis* that using proximity labeling on proteins at the host-pathogen interface captures novel *in vivo* protein-protein interactions during development which enables a new modality for understanding how chlamydiae survive in their intracellular niche^19^.

We drew inspiration from methods previously established for *Streptococcus pyogenes* where the outer membrane proteome was studied by transiently adding proteases to axenic cultures of live bacteria to liberate peptides derived from outer membrane proteins^20^. We hypothesized that these methods could be adapted for virulent Mtb to define the outer membrane proteome, potentially revealing antigenic targets and additional proteins that mediate host-Mtb interactions. We optimized this protease shaving protocol for mycobacteria and applied it to the Mtb strain, H37Rv, revealing novel constituents of the Mtb outer membrane proteome as well as leveraging these basic biologic discoveries to enable new biotechnologies for understanding host-pathogen interactions during Mtb infection. Our results help illuminate potential novel antigenic targets as well as suggest novel mechanisms by which existing Mtb vaccines currently in phase 3 clinical trials may mediate protection. Our study also begins to elucidate rules of functionalizing the surface of Mtb through epitope tagging of one of the surface proteins. We showcase the viability of using this functionalization to chemically conjugate cargo to the surface of Mtb and enable new biological applications that will help facilitate an improved understanding of Mtb-host interactions.

## Results

### Protease shaving can be applied for study of virulent Mtb

A major gap in our understanding of Mtb is the localization of Mtb proteins, especially of those proteins that are accessible on the outer membrane. Addressing this gap in knowledge would facilitate the rational design of antibodies that can target intact Mtb or better identify Mtb proteins that can interact with host machinery (Fig. 1A). Previous studies have successfully identified and validated Mtb outer membrane proteins using targeted approaches. Mitra and colleagues tagged specific Mtb proteins with epitope tags and subsequently performed ultracentrifugation to determine if tagged Mtb proteins were in fractions associated with the outer membrane^21^. Wang and colleagues leveraged bacterial genetics to recover a PPE51 mutant that was resistant to 3bMP1, a novel inhibitor of Mtb growth and survival. Subsequent biochemical analysis using an epitope-tagged PPE51 combined with ultracentrifugation or flow cytometry revealed that PPE51 was an Mtb outer membrane protein^14^. A key similarity between these studies is that they were both targeted methods that first relied on tagging an Mtb protein of interest and then using ultracentrifugation and flow cytometry to confirm protein localization.

**Figure 1:**
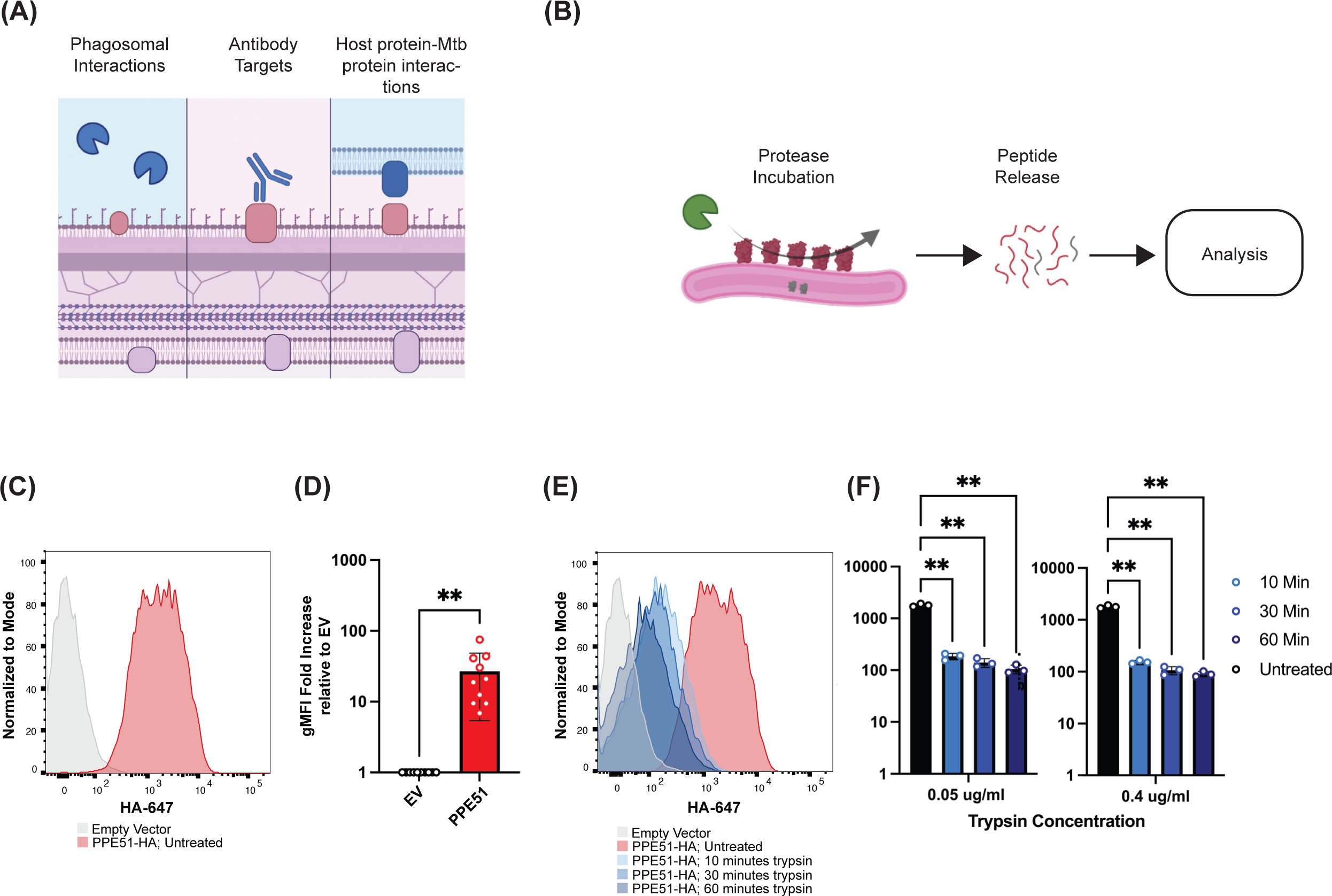
Protease shaving enables the identification of Mtb outer membrane proteins. (A) Schematic of the Mtb outer membrane and potential points of interaction between the Mtb outer membrane and the host immune response. (B) Protease shaving methodology. Intact live Mtb are exposed to protease for defined periods of time and peptides liberated into the supernatant are identified and/or quantified by mass spectrometry. (C) Flow cytometry analysis of HA surface expression on an H37Rv strain expressing an empty vector PPE51-HA. (D) Quantification of HA surface expression on empty vector or PPE51-HA H37Rv strain (** p<0.01, paired t-test). (E) Flow cytometry analysis reveals loss of PPE51-HA signal following trypsin (0.4 μg/ml) exposure. (F) Quantification of HA surface expression on PPE51-HA expressing Mtb over time at two different trypsin (0.4 μg/ml, 0.05 μg/ml) (** p<0.01; two-way ANOVA with Tukey’s multiple comparisons test).

We drew inspiration from prior work in *S. pyogenes* which established protease shaving as a technique to define the outer membrane proteome of bacteria in a proteome-wide manner. In protease shaving, intact and live bacteria are transiently incubated with a protease to liberate protease accessible peptides from the bacterial surface (Fig. 1B)^20^. Liberated peptides are then identified using liquid chromatography and tandem mass spectrometry. Despite the potential ease of use of this technique, to date, it has not been systematically utilized in the study of the mycobacterial outer membrane proteome.

We first sought to test the feasibility of utilizing protease shaving to determine the Mtb outer membrane proteome. To establish feasibility as well as optimize our protocol, we leveraged the previously identified Mtb outer membrane protein, PPE51, to define the biochemical parameters that govern successful protease shaving in Mtb. We generated an H37Rv strain expressing PPE51 with a C-terminal HA tag. Liberation of the HA tag from PPE51 would represent successful shaving of peptides from the Mtb surface, and can be easily read out via flow cytometry using an anti-HA antibody. We first utilized indirect flow cytometry using an anti-HA antibody to validate that PPE51 was antibody accessible without fixation or permeabilization of the bacteria. Compared to an empty vector, fluorescent antibody signal was significantly higher in Mtb expressing PPE51-HA (Fig. 1C-D). After confirming that the HA tag on PPE51 was surface accessible, we next sought to identify protease incubation times and concentrations that resulted in loss of HA staining on the Mtb surface. We incubated Mtb in a sucrose buffer with two different concentrations of trypsin (0.4 μg/mL and 0.05 μg/mL, Methods) and across three different incubation times (10, 30, and 60 minutes). For both trypsin concentrations, we saw a reduction in HA signal as early as 10 minutes. At all timepoints, both protease concentrations resulted in a statistically significant decrease in HA signal as quantified by flow cytometry (Fig. 1E-F). Together, these results suggested that exposure to trypsin can digest Mtb outer membrane proteins and cleave Mtb-derived peptides, thus confirming the utility of protease shaving to define the outer membrane proteome of Mtb.

### Quantitative protease shaving reveals the outer membrane proteome of Mtb

Our flow cytometry data show that protease shaving liberates Mtb peptides from the cell surface. Given this success, we next sought to use this technique to identify novel proteins in the Mtb outer membrane proteome. Based on our results with PPE51, we decided to perform all experiments with a 30-minute incubation with trypsin. To distinguish tryptic peptides cleaved from true outer membrane-tethered proteins from peptides derived from proteins in the culture supernatant, we used trypsinized cell-free Mtb culture supernatant, which would contain proteins secreted by Mtb or proteins liberated because of autolysis, as a control.

We next utilized quantitative mass spectrometry to quantify peptides across these two experimental conditions at the two different trypsin concentrations as determined by our PPE51 pilot experiments. We grew H37Rv Mtb to late log phase and then performed trypsin incubation with either cell-free supernatant or intact, live Mtb. Following protease incubation, peptides from either condition were isolated and labeled using isobaric mass tags for quantitative mass spectrometry. Multiple biological replicates were pooled into a single quantitative mass spectrometry analysis to maximize peptide identification and overlap. Several proteins were identified across both trypsin concentrations that have an elevated abundance in the samples derived from trypsin incubation with intact Mtb compared to cell-free supernatant (Fig. 2A-B, Supplemental Tables 1-2). In the 0.4 μg/mL trypsin conditions, proteins with a fold change > 2 in the intact Mtb condition included a variety of protein types. For example, several hits have unknown functions but are thought to be involved in lipid and fatty acid processes while many of the other proteins are from the type VII secretion system (T7SS), specifically from the PE/PPE protein family. PPE38 was one of the identified proteins and is essential to the secretion of two subsets of ESX-5 type VII secretion system substrates, PPE-PGRS and PPE-MPTR proteins ^22^. Previous work in a PPE38 mutant of *Mycobacterium marinum* found that this protein was exposed on the cell surface^23^. Furthermore, we also recovered PPE51 which we had used in our optimization experiments. In the 0.05 μg/mL trypsin conditions, proteins with a fold change > 2 in the intact Mtb conditions included a similar subset of proteins found in the higher trypsin concentration. One of the proteins found was LprO. Although this particular lipoprotein does not have a known function, the lipoproteins as a family are often involved in immunoregulatory processes and virulence, and some promote an immune response^24^. A hit found across both concentrations, PPE60, is known to be surface-exposed in BCG and has the ability to activate dendritic cells in a TLR2–dependent manner to induce Th1 and Th17 immune responses^25^. This suggested that our protein shaving protocol was identifying genuine Mtb outer membrane proteins.

**Figure 2:**
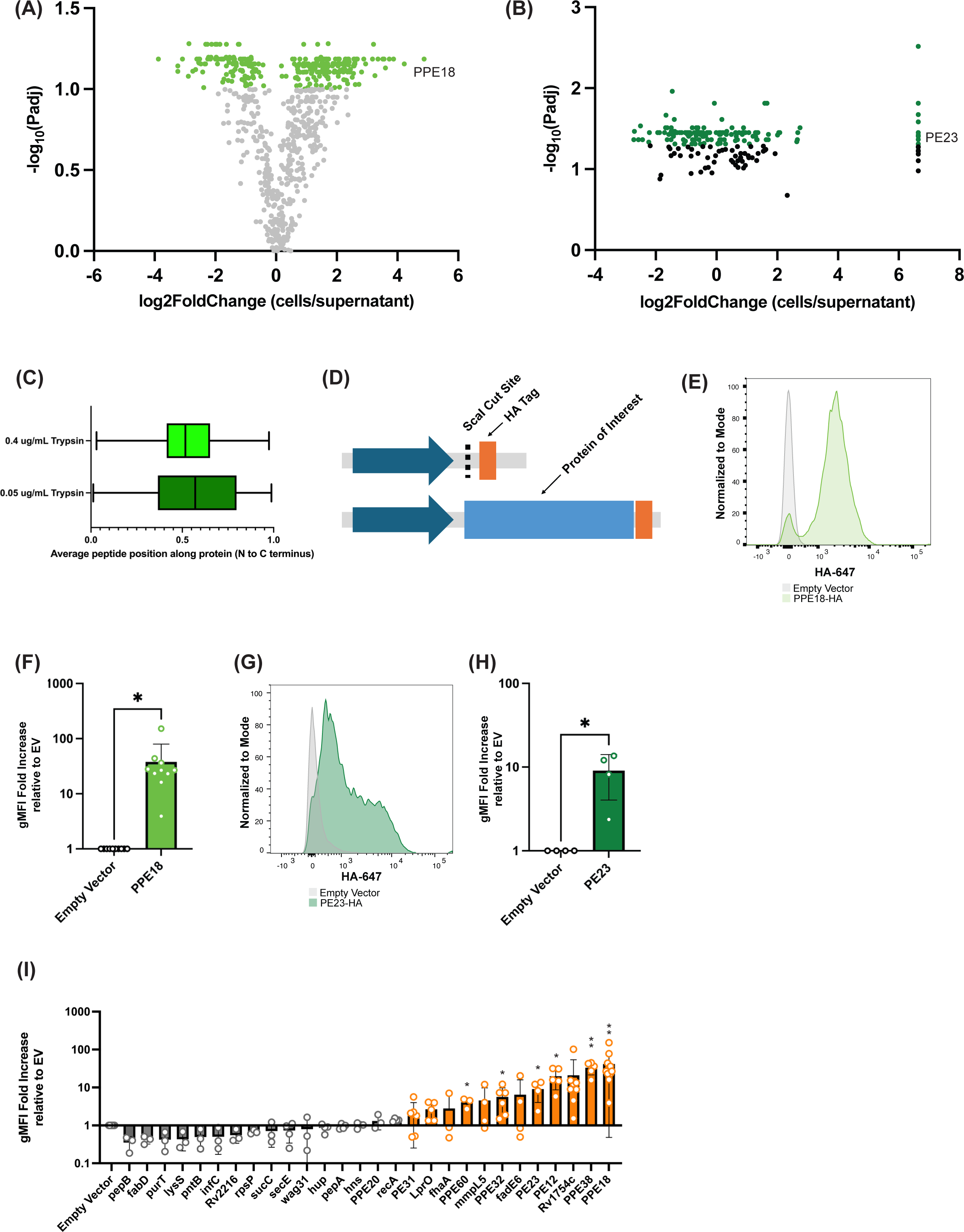
Quantitative mass spectrometry analysis of protease shaving experiments and validation of results. (A) Volcano plot of protease shaving results (trypsin concentration = 0.4 μg/mL). The log FC indicates the abundance ratio of all proteins identified (trypsinized Mtb/trypsinized supernatant). Each point represents one protein. Light green dots indicate proteins with positive fold change of greater than 2, with adjusted p-value of 0.1 or less. (B) Volcano plot of protease shaving results (trypsin concentration = 0.05 μg/mL). The log FC indicates the abundance ratio of all proteins identified (trypsinized Mtb/trypsinized supernatant). Each point represents one protein. Dark green dots indicate proteins with positive fold change of greater than 2, with adjusted p-value of 0.1 or less. (C) Violin plot of the average position of protease shaving results across both trypsin concentrations. The plots show the average position of peptides along proteins that met the statistical significance and >2 positive fold change increase criteria. (D) Plasmid design of surface display vector for validation of protease shaving mass spectrometry results. Each Mtb protein was cloned downstream of a GroEL2 promoter and upstream of a glycine-serine linker and HA tag. (E) Flow cytometry analysis of PPE18-HA, top hit from 0.4 μg/mL protease shaving experiment compared to an empty vector. (F) Quantification of gMFI of HA signal across PPE18-HA and empty vector Mtb strains (* p<0.05, paired t-test). (G) Flow cytometry analysis of PE23-HA, top hit from 0.05 μg/mL protease shaving experiment compared to an empty vector. (H) Quantification of gMFI of HA signal across PE23-HA and empty vector Mtb strains (* p<0.05, paired t-test). (I) Summary of results across all validation strains constructed. Mtb strains with greater than a 2-fold change relative to the empty vector are orange (* p<0.05, ** p<0.01; paired t-test between POI and EV).

We sought to orthogonally validate the protein localization predictions of our protease shaving experiments. We reasoned that we could express epitope tagged variants of Mtb proteins in H37Rv and then use flow cytometry to validate their localization as we did with PPE51. We first conducted a N- vs. C-terminal bias analysis on the Mtb peptides identified via protease shaving to inform our epitope tagging strategy. We reasoned that quantifying this positional bias would facilitate a generalizable cloning strategy where a common vector backbone could be designed including an epitope tag that the candidate outer membrane protein could subsequently be subcloned in. For each Mtb protein identified that met the significance and fold change criteria, the average relative position of the identified peptides along the protein from whence it came was quantified. Proteins with peptides largely originating from the N-terminus of the protein would have an average relative position value closer to 0, whereas peptides originating from the C-terminus would have a value closer to 1. The average relative peptide position for the high and low trypsin concentration groups were 0.53 and 0.57, respectively (Fig. 2C) which did not clearly identify an optimal N- or C-terminal tagging strategy, so we therefore opted with a C-terminal tagging strategy given prior work tagging Mtb outer membrane proteins^21^. We thus proceeded with C-terminal tagging for all candidate proteins. We generated a Gibson Assembly-compatible episomal vector consisting of a strong promoter (*GroEL2*) followed by a ScaI cut site, a short linker and an HA tag (Fig. 2D). To validate the results of our protease shaving, we initially generated two plasmids based expressing a top hit from either the high or the low trypsin concentration experiments expressing PPE18 or PE23, respectively. These plasmids were transformed into H37Rv and then grown in axenic culture prior to flow cytometry analysis. Indirect flow cytometry confirmed that these proteins are surface accessible (Fig. 2E-H). Together, these results indicate the feasibility of identifying outer membrane proteins using quantitative protease shaving.

We next sought to generate additional Mtb strains including additional hits from the protease shaving screen using the identical strategy used for PPE18 and PE23. The top 10 and 25 proteins from the 0.4 μg/mL trypsin and 0.05 μg/mL trypsin concentrations, respectively, were selected for strain generation. Since three proteins overlapped between the two protease concentrations, a total of 32 plasmids were constructed (Supplemental Table 3). All 32 plasmids were transformed into H37Rv. We successfully recovered colonies for all transformations except for the espA overexpressing strain. We grew all the Mtb strains to late log phase and then performed flow cytometry. In each experiment, an empty vector control was stained as a negative control. For each strain, the geometric mean fluorescence intensity (gMFI) of the antibody signal was calculated and then normalized relative to the empty vector. Of the 31 strains generated, 12 Mtb strains showed a positive fold change relative to the empty vector (Fig. 2I). Of those, six had statistically significant fold changes: PPE60, PPE32, PE23, PE12, PPE38 and PPE18. PPE18 consistently had the highest statistically significant fold increase, at an average of 41.63 gMFI-fold change.

We next sought to confirm that our failure to validate surface localization in certain strains was due to the protein being confined to the cytosol rather than simply failing to express. We first confirmed the expression of a subset of these proteins by western blotting for the HA tag. We detected robust expression of recA, hns, and secE proteins by western blotting at the correct molecular weights despite not detecting them by flow cytometry (Sup. Fig. 1). The hup protein fusion could not be detected by western blotting suggesting that its overexpression is not well-tolerated by H37Rv. We next reasoned that lack of antibody staining could be due to the C-terminal HA tag being embedded in the outer membrane while the N-terminus is surface exposed. Therefore, we examined if the successfully expressed protein fusions whose localization could not be validated by flow cytometry had a N-terminal bias in the peptides detected by protease shaving and we did not see an obvious pattern to the proteins that could not be validated. We conclude that it is critically important to use orthogonal validation techniques to confirm the results of protease shaving.

### Large protein fusions to PPE18 fail to export to the Mtb outer membrane

We next sought to confirm that the observations made with PPE18 using an HA tagging strategy are generalizable to other epitope tags. We generated plasmids expressing PPE18 with C-terminal Myc (10 amino acids, 1.2 kDa), V5 (14 amino acids, 1.42 kDa), and 3xFLAG (24 amino acids, 2.73 kDa) tags. We subsequently transformed these plasmids into H37Rv. Indirect flow cytometry confirmed that all epitope tags tested are surface accessible (Fig. 3A-F).

**Figure 3:**
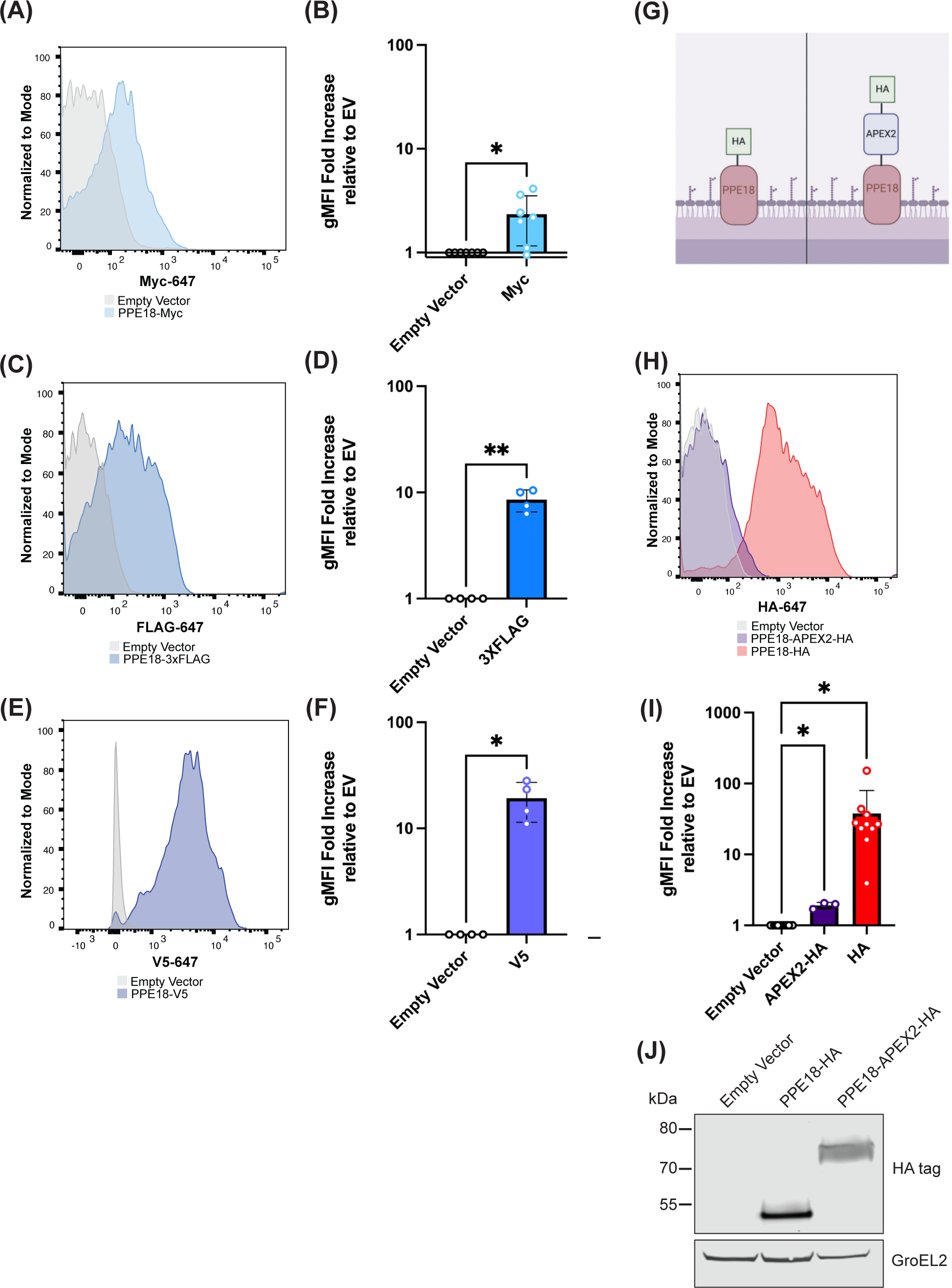
Small epitope tag fusions to PPE18, but not larger protein fusions, are robustly antibody accessible on Mtb. PPE18 fusions were made to Myc, 3xFLAG, and V5 and validated by flow cytometry. Quantification is shown relative to an empty vector (A-B: Myc, C-D: 3xFLAG, E-F: V5, * p<0.05, ** p<0.01; paired t-test). (G) Diagram of PPE18-APEX2-HA fusion. (H) Flow cytometry analysis of HA expression on Mtb expressing PPE18-HA, PPE18-APEX2-HA or an empty vector. (H) PPE18-APEX2 validated for surface localization through HA tagging and measurement of HA surface expression via flow cytometry. (I) Relative quantification is normalized to the empty vector strain (* p<0.05, ** p<0.01; paired t-test). (J) Western blot analysis of HA expression in proteins extracted from pellets of Mtb expressing PPE18-HA, PPE18-APEX2-HA or an empty vector. An HA antibody was used to detect the tagged protein and a GroEL2 antibody was utilized as a loading control.

Given the success of adding epitope tags to PPE18, we next asked whether larger proteins could also be exported to Mtb’s outer membrane. Several biotechnologies desired in the field would be enabled with an Mtb strain that can be functionalized with heterologous proteins including surface display of heterologous enzymes to monitor bacterial infection, phagocyte interaction, and characterization of pathogen-specific compartments. Previous studies have shown that APEX2, an enzyme that facilitates proximity labeling, could be targeted to the periplasm of Mtb and *Mycobacterium smegmatis* ^26,27^. We thus sought to determine if fusion of APEX2 containing a C-terminal HA tag to PPE18 (PPE18-APEX2-HA) would facilitate export of APEX2 to the Mtb surface (Fig. 3G). Compared to an Mtb strain expressing PPE18-HA, we observed far less HA staining of the PPE18-APEX2-HA strain (Fig. 3H-I). This could not be explained by poor expression of the construct because western blot analysis of the HA-tagged protein revealed robust expression by Mtb (Fig. 3J). We thus conclude that while small epitope tags can be delivered to the Mtb surface by fusion to PPE18, larger proteins cannot be. Similar conclusions were made using PPE18 with another proximity labeling enzyme, TurboID (Sup. Fig. 2).

### PPE18-ALFA tag protein fusions support investigation of PPE18 localization and surface functionalization of Mtb

Given that PPE18 is an antigen included in the M72 vaccine, which is presently in vaccine trials, we sought to obtain an enhanced understanding of PPE18 localization on the Mtb surface using microscopy. While flow cytometry demonstrates the surface accessibility of PPE18, it cannot evaluate the spatial distribution of proteins along the bacterium^28^.

We generated an additional Mtb strain where PPE18 was tagged with an ALFA tag (1.69 kDa), a rationally designed epitope tag for which a high affinity nanobody reagent has been developed ^29^. Using flow cytometry, we first confirmed that the ALFA tag was surface accessible (Fig. 4A-C). We utilized image cytometry to orthogonally confirm the spatial distribution of PPE18 on Mtb. Mtb expressing an empty vector or PPE18-ALFA were grown in the presence of HADA, a blue, fluorescent D-amino acid that labels Mtb cell wall peptidoglycans. Following HADA incorporation, cells were directly stained with an anti-ALFA nanobody conjugated to Alexa Fluor 647. We observed robust HADA incorporation in both empty vector and PPE18-ALFA strains. Among the cells that were HADA positive, the percentage of Mtb that were AF647 positive was significantly greater in the PPE18-ALFA strain than the empty vector control (Fig. 4D-E). The resolution of image cytometry was insufficient to determine if there are specific spatial distribution patterns of PPE18-ALFA on the Mtb surface, so we next turned to confocal microscopy. Imaging the PPE18-ALFA strain grown axenically revealed a punctate distribution of PPE18 along the body of the bacterium, representing to our knowledge the first visual inspection of Mtb surface protein spatial distribution (Fig. 4F). Similar findings were obtained when we imaged the PPE18-HA strain by confocal microscopy (Sup. Fig. 3).

**Figure 4:**
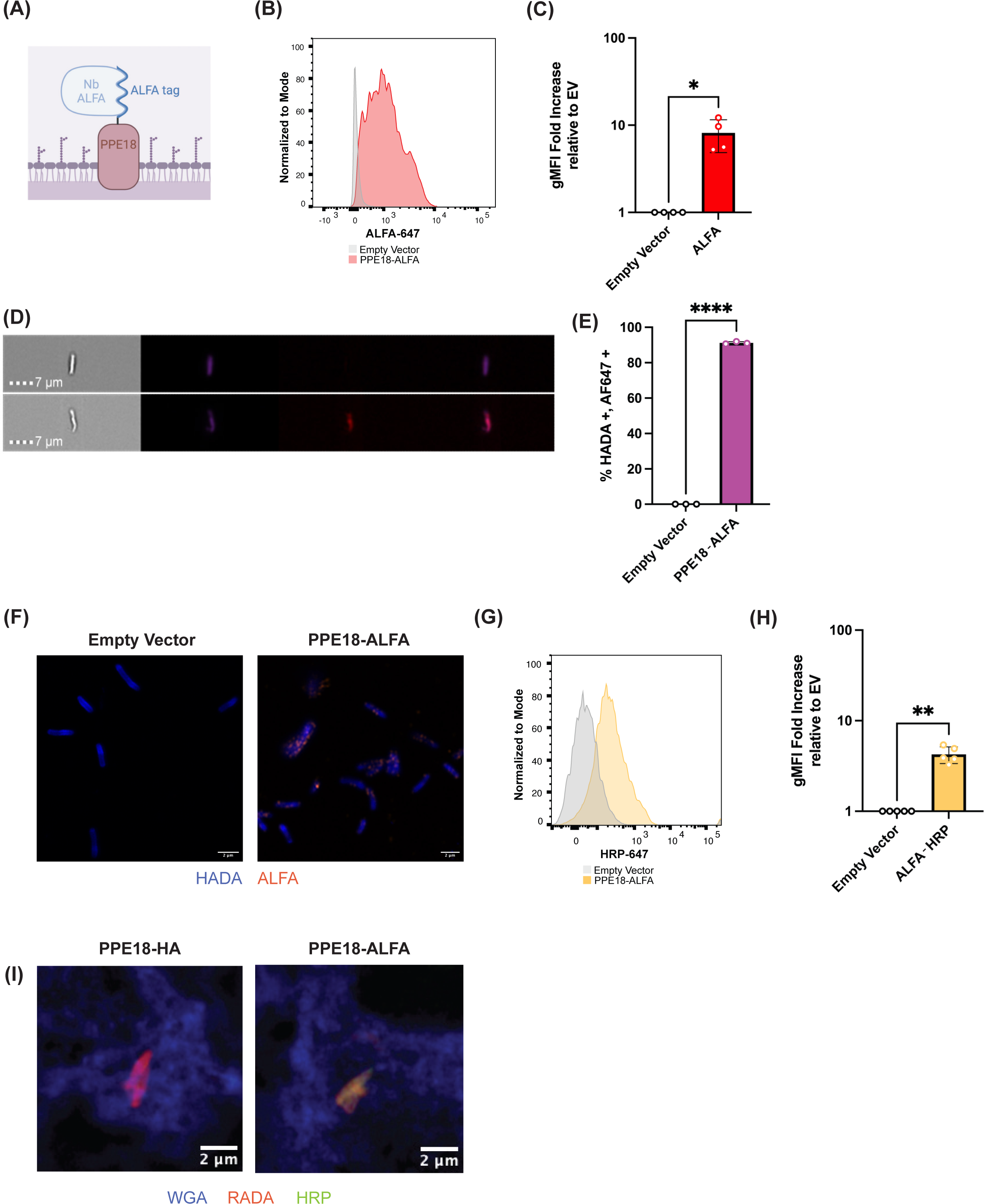
PPE18-ALFA fusions can be used to study protein localization and facilitate delivery of proteins to Mtb-containing phagosomes. (A) Diagram of PPE18-ALFA fusion with bound anti-ALFA nanobody. (B) A PPE18 fusion was made to the ALFA tag, and flow cytometry was conducted using an anti-ALFA nanobody conjugated to AlexaFluor 647. (C) Quantification of gMFI of ALFA signal across PPE18-ALFA and empty vector Mtb strains (* p<0.05, paired t-test). (D) Image cytometry analysis of anti-ALFA AlexaFluor 647 on HADA stained Mtb (empty vector or PPE18-ALFA). (E) Quantification of image cytometry analysis of anti-ALFA AlexaFluor 647 staining of Mtb expressing either an empty vector or PPE18-ALFA (**** p<0.0001, paired t-test). (F) Representative images of anti-ALFA staining of Mtb expressing either an empty vector or PPE18-ALFA using Airyscan microscopy (50x). Both Mtb strains were grown with HADA to visualize their cell walls (blue) and were stained for anti-ALFA AlexaFluor 647 (orange). Scale bar indicates 2 µm. (G) Mtb strains expressing an empty vector or PPE18-ALFA were subject to surface conjugation with an anti-ALFA nanobody fusion to HRP. Following surface conjugation, indirect flow cytometry was performed to detect the presence of HRP on the Mtb surface. (H) Quantification of gMFI of HRP signal across PPE18-ALFA and empty vector Mtb strains following conjugation to an anti-ALFA nanobody fusion to HRP (** p<0.01, paired t-test). (I) Microscopy analysis of surface engineered Mtb in macrophages. PPE18-HA or PPE18-ALFA Mtb strains were incubated with an ALFA tag nanobody fused to HRP and delivered to macrophages. Following phagocytosis, macrophages were fixed and stained for macrophage membranes (WGA-405) and HRP (AF488) (blue = macrophage membranes, red = Mtb cell wall, green = HRP). Shown is the merged image (see supplemental figure 4, for split channels). Scale bar indicates 2 µm.

Based on our success with display of the ALFA tag on the Mtb surface and our lack of success in creating Mtb protein fusions to PPE18 that result in outer membrane delivery of large heterologous proteins, we reasoned that an alternative strategy to leverage the outer membrane proteome to create Mtb strains with novel functions on their surface would be to leverage the exquisite affinity of the anti-ALFA nanobody and fuse the nanobody to diverse proteins of interest. An approach like this is modular and can be improved upon as more affinity reagents are developed for native Mtb proteins and more epitope tags with companion affinity reagents are engineered. To demonstrate this principle, we utilized the PPE18-ALFA strain and functionalized the Mtb surface with horseradish peroxidase (HRP), an enzyme capable of performing proximity labeling reactions ^30,31^. We grew Mtb expressing PPE18-ALFA or an empty vector and then incubated them with an anti-ALFA nanobody-HRP fusion. We next confirmed conjugation of HRP to the Mtb surface using indirect flow cytometry. Compared to the empty vector Mtb strain, HRP was much more abundant on the PPE18-ALFA surface (Fig. 4G-H).

We lastly sought to leverage our engineered Mtb in an application for host-pathogen interactions and deliver surface-engineered Mtb to macrophages. Mtb expressing an PPE18-HA or PPE18-ALFA were grown in the presence of RADA, an orange-red fluorescent D-amino acid that labels Mtb cell wall peptidoglycans. Following RADA incorporation, Mtb were directly stained with the ALFA nanobody-HRP fusion and used to infect primary human monocyte-derived macrophages at a multiplicity of infection of 1. After infection, the macrophages were fixed, stained with wheat germ agglutinin 405 (WGA-405) to visualize host cell membranes and prepared for immunofluorescence detection of HRP protein using primary and secondary antibody staining. Airyscan confocal microscopy revealed colocalization between RADA and HRP in our PPE18-ALFA Mtb strain which was absent in the PPE18-HA strain (Fig. 4I, Sup. Fig. 4). This demonstrates that surface functionalized Mtb can be delivered to a dynamic, complex setting like the host phagosome.

## Discussion

Despite the importance of the mycobacterial outer membrane to the regulation of Mtb physiology in axenic culture as well as in phagocytes, much of our understanding of the mycobacterial outer membrane proteome remains incomplete. Prior studies in Mtb have identified outer membrane proteins, but the techniques deployed to define these proteins focused on individual proteins and used laborious techniques such as ultracentrifugation to confirm their localization. Here we build from approaches originally established in *Streptococcus pyogenes*, to apply protease shaving to live, virulent Mtb. Using these approaches, we nominate and identify novel Mtb outer membrane proteins including an Mtb vaccine antigen, providing insight into the mechanism by which immunity following this vaccine might occur. We further demonstrate that defining the outer membrane proteome of Mtb can be utilized to deliver heterologous proteins to the Mtb surface.

Several of the top proteins identified in protease shaving that were validated shared notable biological characteristics. Namely, many of these proteins belong to the PE and PPE family, known for their proline and glutamic acid residue-rich N-terminal sequences. The PE/PPE proteins are hypothesized to have functions like nutrient transport, host-pathogen interaction, or immune evasion - functions that necessitate either their secretion or cell wall or outer membrane surface localization^32^. While the subcellular localization of PE/PPE family members remains incompletely defined, our findings suggest that many members are surface exposed. We saw both known outer membrane localized hits and expanded the list of proteins that are known to be surface accessible. Examples of known outer membrane localized proteins that we also confirmed in this study include: PPE60, PE12, PPE32, and PPE38. For example, Su and colleagues demonstrated that PPE60 was cell surface exposed through a recombinant BCG PPE60-His strain that underwent immunoblotting and a proteinase K degradation assay^25^. Their study, along with another by Gong et al, highlight the pro-inflammatory immunological role of PPE60 which includes the promotion of dendritic cell maturation and macrophage pyroptosis ^33^. Similarly, PE12, PPE32, and PPE38 were all found to be surface associated and found to have some effect on immune response such as inhibiting the transcription of inflammatory cytokines^34^, promoting ER stress mediated apoptosis ^35,36^, and downregulating MHC-I antigen presentation ^23,37^, respectively. In contrast, very little is known about the localization and function of PE23. Previous studies did not identify it as a likely outer membrane protein and to our knowledge, we are the first to showcase its surface accessibility ^18,38^.

Of the 32 proteins tested, PPE18 was of particular interest. Although not previously shown to be on the Mtb surface, other studies have identified PPE18 by mass spectrometry in Triton X-114 extracts, cell membrane protein fraction and whole cell lysates from Mtb H37Rv ^39,40^. We demonstrated surface localization of this protein and found that it consistently had the highest fold change increase of geometric mean fluorescence intensity (gMFI) when compared to our empty vector control and all other proteins in this study. Moreover, we were able to show direct evidence of PPE18’s surface localization and spatial organization via microscopy. PPE18’s localization likely underscores its function. For instance, previous studies have suggested that PPE18 is a virulence factor that plays a role in IL-10 production of macrophages by interacting with TLR2 ^41,42^. PPE18 is also one of the proteins included in an Mtb vaccine, M72/AS01E, and was originally chosen because of its ability to stimulate cell-mediated and antibody-mediated immune responses in both mice and controlled human infection ^43^. Specifically, PPE18 is one of a handful of PE/PPE proteins known to generate CD4 mediated T-cell responses which was showcased in the high frequency of M72-specific CD4+ T cells after dosage^44,45^. Additionally, antibodies are generated against PPE18 following vaccination^46^. Antibodies against Mtb antigens have steadily gained research interest due to their ability to be elicited during vaccination ^47,48^, their ability to be used as diagnostics^49^ , and role in mediating antimicrobial responses ^50–52^. Altogether, this further supports the ability of surface shaving to identify outer membrane proteins of relevant clinical and therapeutic interest.

In addition to identifying surface proteins, we were able to discern several characteristics of surface functionalization for Mtb. One is the ability to localize small peptides, but not larger proteins, via fusion to surface-accessible Mtb proteins. Our ability to fuse five different epitope tags to PPE18 is in agreement with previous studies that have used similarly sized tags to detect the expression of recombinant proteins in mycobacteria ^53^. For larger proteins, previous strategies were devised that fuse signal peptides from validated substrates of the Sec protein secretion system in order to dictate localization of an enzyme to a compartment of interest, like the periplasm^27^. This strategy is less optimal for purposes studying the Mtb surface interface as many of the surface accessible, outer membrane proteins we identified were associated with the ESX systems of the Type VII secretion system, where a verified signal peptide is not known. To anchor an enzyme of interest to the Mtb surface, we hypothesized that fusing it to a known outer membrane protein might be one way to facilitate its export. We were not successful at achieving this goal, and several outstanding questions still need to be answered including the upper limit of protein size to be fused, whether additional forms of optimization need to take place such as modulating protein expression level, and the underlying biological mechanisms of export to the Mtb surface. Nevertheless, our work provides a framework to display short binding motifs on the surface of Mtb.

Lastly, we successfully leveraged the ALFA tag fused to PPE18 to functionalize the Mtb surface with the enzyme HRP. We showed that the ALFA tag nanobody-HRP fusion was more abundant on the Mtb surface than our empty vector nanobody-HRP fusion control in an axenic culture. HRP is a peroxidase of interest because it is able to convert substrates like biotin-phenol, in the presence of H₂O₂, into a reactive radical that covalently tags electron-rich amino acids of neighboring proteins (proximity labeling)^54^. Moreover, HRP is reactive in oxidizing environments like the extracellular regions and can be used to proteomically map the surface of living cells^30^. This system could aid in the identification of endogenous host or Mtb interaction partners of surface proteins or the spatial resolution and composition of surface proteins and how they change given mycobacterial phase of growth. Our data also show that we can deliver these modified bacteria to the phagosome. Previously used techniques to study the Mtb-containing phagosome include mechanical homogenization paired with differential centrifugation or conjugation of magnetic beads onto the pathogen’s surface, both of which have their own disadvantages such as cellular component contamination and artificial altering of phagosomal interactions, respectively ^55,56^. Although several studies have started to characterize the Mtb phagosome with targeted methods, obtaining useful information around its dynamics, these methods focus on one-dimensional profiling of the pathogen-containing phagosome ^57–60^. Consequently, proximity labeling that is enabled by functionalization of the Mtb surface could facilitate the holistic, pure isolation of the Mtb-containing phagosome. Future uses of this technology will enable the compositional analysis of the Mtb-containing phagosome using proximity labeling which will greatly enhance our understanding of Mtb-host interactions. In addition to using the ALFA tag as a conjugation strategy, we further hypothesize that additional tag systems such as the covalent SpyTag-SpyCatcher system could be used to covalently modify the Mtb surface given the convenient size of the SpyTag (1.68 kDa) ^61^. In conclusion, this work further characterizes the Mtb outer membrane proteome and demonstrates that this compartment can be exploited to identify new vaccine targets and set the stage to enable biotechnologies for understanding pathogen interactions.

## Materials and Methods

### Mtb culture for protease shaving and flow cytometry

Mtb H37Rv was grown in 7H9 media supplemented with 10% OADC (Thermo, B12351), 0.2% glycerol (Thermo, 17904), and 0.02% tyloxapol (VWR, TS42237-0050), to mid-log phase (7H9-Tyloxapol).

### Plasmid Construction and Mtb Transformation

For validation of protease shaving mass spectrometry results, full-length Mtb genes (without a C-terminal stop codon) were amplified from H37Rv genomic DNA. PCR amplicons were inserted by Gibson Assembly into a standard surface display vector consisting of a strong promoter (GroEL2), followed by a ScaI cut site for gene insertion, a glycine-serine linker and an HA tag.

Additional PPE18 fusion strains (Myc, 3xFLAG, V5, ALFA, APEX2-HA, TurboID-HA) were generated by Gibson assembly.

### Mtb culture for transformations and electroporation

Mtb H37Rv was grown in 7H9 media supplemented with 10% OADC (Thermo, B12351), 0.2% glycerol (Thermo, 17904), and 0.02% Tween-80 (Sigma-Aldrich, P1754-500ML), to mid-log phase (7H9-OADC).

Wildtype H37Rv was grown in 7H9 to mid-log phase. Bacteria were pelleted and washed with 10% glycerol three times and then resuspended in 400 μL of 10% glycerol per transformation. They were then transformed with 200 ng of plasmid via electroporation. Electroporated bacteria were recovered in 1 mL 7H9 for 48 hours. They were plated on 7H10 plates supplemented with 50 μg/mL hygromycin B (Millipore Sigma, 10843555001) for 17 days. Transformants were inoculated into 5 mL of 7H9 containing hygromycin B for 5 days prior to subsequent dilution and storage at -80°C.

### Mtb staining, surface conjugation and flow cytometry

Mtb strains were grown up in 7H9 with 50 μg/mL hygromycin B to an OD600 between 0.5 and 1. 1e8 bacteria from each strain were pelleted and resuspended in 500 μL 4% paraformaldehyde (VWR, AAJ61899-AP) for 1 hour at room temperature. Following fixation, Mtb were pelleted for 5 minutes at 4000xg and washed with 500 μL FACS buffer (PBS without divalent cations, 2% heat-inactivated fetal bovine serum, 2 mM EDTA). Bacteria were incubated with the indicated primary antibody (1:250) which included rabbit anti-HA (CST, C29F4 #3724), chicken anti-myc (exalpha, ACMYC), rabbit anti-3xFLAG (CST, D6W5B #14793), rabbit anti-V5 (CST, D3H8Q #13202), in FACS buffer for 30 minutes at room temperature. Alexa Fluor 647-conjugated secondary antibodies were added according to the species of the primary antibody (Thermo, A-21245, rabbit or Thermo, A-21449, chicken) at a 1:120 dilution for an additional 10 minutes in the dark. The bacteria were washed two times with FACS buffer, resuspended in 500 μL of FACS buffer and analyzed by flow cytometry.

In the case of the ALFA tag, bacteria were stained directly with the FluoTag-X2 anti-ALFA AF647 (NanoTag, N1502) (1:500) for an hour, washed two times with FACS buffer, and resuspended in 500 μL FACS buffer prior to analysis by flow cytometry.

For surface conjugation of HRP to the Mtb surface, PPE18-HA or PPE18-ALFA expressing bacteria were stained with sdAB anti-ALFA HRP (NanoTag, N1505) (1:500 in FACS buffer) for an hour at room temperature. Bacteria were subsequently stained with an anti-HRP antibody (Thermo, MA1-10371) (1:250) in FACS buffer for 30 minutes at room temperature. Mtb were finally stained with an AF647-conjugated secondary antibody (1:120) for 10 minutes at room temperature prior to two washes with FACs buffer and analysis by flow cytometry.

### Protein Shaving Optimization Experiments

Surface proteins of Mtb H37Rv were identified using a protocol adapted from prior literature ^20^. 10 mL of H37Rv expressing PPE51-HA were grown in 7H9-Tyloxapol to mid-log phase. 1e8 bacteria from each replicate per condition were pelleted 3000 x g for 5 minutes, washed once in PBS without divalent cations (Corning, 21-040-CV), and resuspended in a 100 μL 40% sucrose/PBS buffer. For experimental conditions where Mtb were exposed to protease, either 0.4 μg/mL or 0.05 μg/mL of trypsin (Promega, V5113) was added to the sucrose buffer. For the negative control condition, PBS without divalent cations was added. Samples with protease were incubated for 10, 30, or 60 minutes. Following incubation, all samples were then pelleted 3000xg for 5 minutes, washed once in PBS, and resuspended in 500 μL 4% paraformaldehyde for 1 hour at room temperature. Samples were then stained for flow cytometry as described above.

### Protein surface shaving protocol

100 mL of Mtb WT H37Rv were grown in 7H9-Tyloxapol per replicate to mid-log phase. Bacteria were pelleted by centrifugation at 3000xg for 10 minutes at 4° C and resuspended in PBS. Three washes total were performed. Following the third wash, the Mtb pellet was resuspended in a 40% sucrose/PBS buffer at a ratio of 100 μL of buffer for every 100 mL of bacterial culture. For the protease shaving experimental conditions, either 0.4 μg/mL or 0.05 μg/mL of trypsin was added and for control conditions, PBS was added. The suspension was incubated for 30 minutes at 37° C. Samples were pelleted by centrifugation at 3000xg for 10 minutes at 4° C and the supernatant was recovered. The supernatant was filtered with a 0.2 μm pore-size filter (Pall, ODM02C35) twice. For the background control conditions, either 0.4 μg/mL or 0.05 μg/mL of trypsin was added to the filtered supernatant and incubated for 30 minutes at 37° C. Oasis HLB Extraction Cartridges (Waters, WAT094225) were used to clean the supernatants for mass spectrometry preparation. The cartridges were equilibrated with 0.6mL of 80% acetonitrile (Sigma-Aldrich, 34851-1L) and subsequently 0.6 mL of 0.1% formic acid (Sigma-Aldrich, 33015). Samples were loaded at a ratio of 300 μL of supernatant to 300 μL of PBS. Each sample was washed twice with 0.6 mL of 2% acetonitrile/0.1% formic acid. Elution of the samples was done in three steps with 0.6 mL of the following solutions: 10% acetonitrile/0.1% formic acid, 20% acetonitrile/0.1% formic acid, and 50% acetonitrile/0.1% formic acid, each collected in its own tube. All samples were snap-frozen in liquid nitrogen and lyophilized.

### Tandem Mass Tag (TMT) labeling

Peptides for mass spectrometry were prepared for TMT analysis according to the manufacturer’s instructions (Thermo, 90061). The tryptic peptides were resuspended in 100 μL of 100 mM TEAB, then vortexed and briefly centrifuged. 41 μL of anhydrous acetonitrile was added to each of the TMT label reagents, they were vortexed and briefly centrifuged. The 41 μL of the TMT reagents were added to 100 μL of sample, vortexed and briefly centrifuged. The samples were incubated for 1 hour at room temperature. After an hour, 8 μL of 5% hydroxylamine were added to each sample and incubated for 15 minutes to quench the reaction. Equal amounts of each sample were then combined and placed into a speed-vac to dry to completion.

### LC-MS-MS and quantitative analysis

The TMT labeled tryptic peptides were resuspended in 10 uL of 0.2% formic acid and 1 μL was injected. The tryptic peptides were separated by reverse phase HPLC (Thermo Ultimate 3000) using a Thermo PepMap RSLC C18 column (2um tip, 75umx50cm PN# ES903) over a gradient before nano-electrospray using a ThermoFisher Orbitrap Exploris 480 mass spectrometer. Solvent A was 0.1% formic acid in water and solvent B was 0.1% formic acid in acetonitrile. The gradient conditions were 1% B (0-10 min at 300nL/min) 1% B (10-15 min, 300 nL/min to 200 nL/min) 1-10% B (15-20 min, 200nL/min), 10-25% B (20-65 min, 200nL/min), 25-36% B (65-75 min, 200nL/min), 36-80% B (75-75.5 min, 200 nL/min), 80% B (75.5-80 min, 200nL/min), 80-1% B (80-80.1 min, 200nL/min), 1% B (80.1-100 min, 200nL/min).

The mass spectrometer was operated in a data-dependent mode. The parameters for the full scan MS were resolution of 120,000 across 375-1600 m/z and maximum IT 25 ms. The full MS scan was followed by MS/MS for as many precursor ions in a two second cycle with a NCE of 36, dynamic exclusion of 30 s and resolution of 30,000.

Raw mass spectral data files (.raw) were searched using Sequest HT in Proteome Discoverer (Thermo). Sequest search parameters were: 10 ppm mass tolerance for precursor ions; 0.05 Da for fragment ion mass tolerance; 2 missed cleavages of trypsin; fixed modification were carbamidomethylation of cysteine and TMT modification on the lysines and peptide N-termini; variable modifications were methionine oxidation, methionine loss at the N-terminus of the protein, acetylation of the N-terminus of the protein and also Met-loss plus acetylation of the protein N-terminus.

Data was searched against a *Mycobacterium tuberculosis* database downloaded from Uniprot and a contaminant database made in house at the Koch Institute Proteomics Core. The search criteria are as follows: FDR for peptide matches of high confidence was set at 0.01, Sequest HT Xcorr greater than or equal to 2.00 and maximum delta mass was set at 3 PPM.

The statistical significance of a protein’s fold change was calculated by performing paired T tests. First, for each replicate, the experimental and control condition abundance levels were divided by the corresponding replicate’s control abundance level. These ratios groups were the inputs to python’s scipy.stats.ttest_rel function, two-sided hypothesis and p-values were obtained. The p-values were then adjusted using python’s statsmodels.stats.multitest.fdrcorrection function for the Benjamini/Hochberg correction. In the cases where the control abundance levels were zero but nonzero in the experimental conditions, values were imputed to be 500 - the lowest raw abundance levels seen, to obtain a fold change ratio. To be considered a potential surface protein, a candidate protein had to have: at least 2 peptides obtained from protease cleavage, greater than 2-fold log change between the experimental (live, intact Mtb) versus control (cell-free supernatant) and have an adjusted p-value of less than 0.1. As all top proteins would be experimentally validated via epitope tagging, the adjusted p-value was chosen as 0.1 to balance the likelihood of the type I error occurring with exploring the search space for potential surface proteins.

### Western blotting

Mtb were first grown in 7H9-Tyloxapol to an OD600 of 0.8-1. Bacteria were pelleted and lysed in RIPA buffer (VWR, 97063-270) with 1X Halt Protease Inhibitors (Thermo, 78430). Bead beating was performed with Lysing Matrix B (MP Biomedical, 16911050-CF) for three rounds of agitation, 45 seconds each. The culture supernatants were 0.2 μm filtered (Pall, ODM02C35) twice. Protein concentration was determined by BCA assay (Thermo) and equal amounts of protein lysate were denatured and prepared as recommended by the manufacturer for western blotting with Bolt 4-12% Bis-Tris gels (Thermo, NW04120BOX). Gels were transferred to nitrocellulose membranes using an iBlot 2 (Thermo , 25-107). Membranes were blocked with Odyssey Blocking Buffer (Licor, 927-60003) and stained with the indicated primary antibodies. They were then stained with secondary antibodies, washed, and imaged on the Licor Odyssey imaging system.

### Immunofluorescence microscopy of axenic culture

Mtb cultures of the empty vector strain or PPE18-ALFA strain were grown to mid-log phase in the presence of HADA (BioTechne , 6647/5) (1:500). 1e8 bacteria from each strain were incubated with anti-ALFA AF647 (Nanotag, N1502) (1:500), washed twice with PBS and resuspended in 500 μL 4% Paraformaldehyde for 1 hour at room temperature. After fixation, both strains were washed once in PBS and 100 μL of each strain resuspended in PBS was pipetted onto a 35 mm dish (Mattek). After evaporation of excess PBS, the strains were covered with a 1% agarose pad. Strains were imaged using a Zeiss CellDiscoverer 7 Microscope at a 50x objective lens.

Mtb cultures of the empty vector strain or PPE18-HA strain were grown to mid-log phase in the presence of HADA (BioTechne, 6647/5) (1:500). 1e8 bacteria from each strain were resuspended in 500 μL 4% paraformaldehyde for 1 hour at room temperature. After washing once with PBS, strains were stained with the rabbit anti-HA primary antibody (CST, C29F4 #3724) (1:250) in FACS buffer for 30 minutes at room temperature. Fluorescent AF647-conjugated goat anti-rabbit (Thermo, A-21245) secondary antibody (1:120) was added and incubated for another 10 minutes in the dark. The bacteria were washed two times with FACS buffer, and 100 μL of each strain resuspended in FACS buffer was pipetted onto a 35 mm dish (Mattek, P35G-1.5-14-C). After evaporation of excess FACS, the strains were covered with a 1% agarose pad. Strains were imaged at 100x magnification using a Zeiss CellDiscoverer 7 Microscope with a 50x objective and 2x tube lens.

### Macrophage Culture

Deidentified buffy coats were purchased from the Massachusetts General Hospital Blood Donor Center. Peripheral blood mononuclear cells (PBMCs) were isolated from buffy coats by density centrifugation. CD14+ monocytes were isolated from PBMCs by positive selection using the EasySep Human CD14 Positive Selection Kit II (Stemcell, 17858). Monocytes were plated in 12 well chamber slides (IBIDI, 81201) phenol-free RPMI media containing heat-inactivated fetal bovine serum (10%), HEPES (10 mM) and L-glutamine (2 mM) supplemented with 25 ng/mL M-CSF (Biolegend, 574806) for 6 days.

### Macrophage Immunofluorescence Microscopy

Slides of Mtb infected macrophages were washed twice with PBS and stained with wheat germ agglutinin (WGA) 405 (Thermo, W56132) (1:400) for 20 minutes. After washing twice with PBS, they were incubated with 0.1% Triton X-100 (Sigma-Aldrich, X100-5ML) for 10 minutes. They were washed three times with PBS and blocked with 3% bovine serum albumin (BSA) (Sigma-Aldrich, A3294) for 1 hour at room temperature. After, slides were stained for 1 hour at room temperature with primary antibody (anti-HRP mouse, Thermo, MA1-10371) diluted in antibody dilution buffer (1% BSA). Wells were washed three times with PBS for 5 min each and stained with Alexa Fluor 647 (AF488)-conjugated goat anti-mouse secondary antibody (Thermo, A32723) diluted 1:5000 in antibody dilution buffer at room temperature for 1 hr. Wells were washed three times with PBS for 5 min, well dividers were removed, and slide covers were mounted on slides using Prolong Diamond antifade mounting media (Thermo, P36970). Slides were imaged using a Zeiss CellDiscoverer 7 Microscope at a 50x objective lens.

### Statistical analyses

Unless otherwise indicated, all statistical tests were performed in GraphPad Prism 10.

## Acknowledgements

We thank the Bryson and Barczak lab members for discussions and feedback. We thank the Koch Institute’s Robert A. Swanson (1969) Biotechnology Center for technical support, specifically the Biopolymers & Proteomics Core.

## Funding

We acknowledge funding support from the Susan and Terry Ragon Foundation (BDB) and the NIH (R35GM142900, R01AI152158, BAA-NIAID-NIHAI201700104). This work was performed in part in the Ragon Institute BSL3 core, which is supported by the NIH-funded Harvard University Center for AIDS Research (P30 AI060354). BAL and SLS were supported by Siebel Foundation Fellowships.

### Author contributions

Conceptualization: BAL, BDB

Methodology: BAL, CRZ, OKL, BLA, SLS, BDB

Investigation: BAL, CRZ, OKL, BLA, SLS, BDB

Data curation: BAL, CRZ, BDB

Formal analysis: BAL, CRZ, BDB

Visualization: BAL, BDB Resources: BDB

Writing - original draft: BAL, BDB

Writing - review and editing: all authors Supervision: BDB

Funding: BDB

## Data and materials availability

All the data supporting this work is included in the figures or can be found in the supplementary information. Mass spectrometry data have been uploaded to the PRIDE Database under accession PXD052936. Plasmids for PPE18-HA and PPE18-ALFA have been deposited to Addgene.

## Supplemental Figure Captions

**Supplemental Figure 1: Western blot analysis of HA protein fusions to Mtb proteins identified by protease shaving.** Mtb expressing an empty vector, PPE18-HA, recA-HA, Rv3852-HA, espI-HA, secE1-HA, and hup-HA were grown to mid-log phase, pelleted, and lysed. Bacterial lysates were probed for the presence of the HA epitope tag and for the GroEL2 protein as a loading control.

**Supplemental Figure 2: HA signal on PPE18-TurboID-HA strains.** (A) Flow cytometry analysis of HA signal on Mtb expressing PPE18-HA, PPE18-TurboID-HA or empty vector stained with an anti-HA antibody. (B) Western blot analysis of HA expression in proteins extracted from pellets of Mtb expressing PPE18-HA, PPE18-TurboID-HA or an empty vector. An HA antibody was used to detect the tagged protein and a GroEL2 antibody was utilized as a loading control.

**Supplemental Figure 3: Microscopy analysis of HA localization on Mtb expressing an empty vector or PPE18-HA.** Representative images of HA localization on Mtb expressing PPE18-HA or an empty vector Mtb strains. Images were obtained using Airyscan microscopy (100x). HADA was used to visualize Mtb cell walls. Scale bar indicates 1 µm for PPE18-HA and 2 µm for the empty vector.

**Supplemental Figure 4: Microscopy analysis of surface engineered Mtb (PPE18-HA or PPE18-ALFA) in macrophages.** PPE18-HA (A) or PPE18-ALFA (B) Mtb strains were incubated with an ALFA tag nanobody fused to HRP and delivered to macrophages. Following phagocytosis, macrophages were fixed and stained for macrophage membranes (WGA-405) and HRP (AF488) (blue = macrophage membranes, red = Mtb cell wall, green = HRP). Shown is the merged image and individual channels. Scale bar indicates 2 µm.

**Supplemental Table 1: Results from protease shaving with 0.4 μg/mL trypsin.**

Quantification of all Mtb proteins detected in mass spectrometry experiments.

**Supplemental Table 2: Results from protease shaving with 0.05 μg/mL trypsin.**

Quantification of all Mtb proteins detected in mass spectrometry experiments.

**Supplemental Table 3: Primer sequences for Mtb protein shaving top hits.** The top 10 and 25 proteins from the 0.4 μg/mL trypsin and 0.05 μg/mL trypsin concentrations, respectively, were selected for strain generation. Three proteins overlapped between the two protease concentrations, thus a total of 32 plasmids were constructed. Primer sequences for the plasmid construction for tagging protein hits with the HA epitope are listed.

## References

1. Becken III, B. A., Rudas, F. J. B. & Chatterjee, A. An update on tuberculosis. in Viral, Parasitic, Bacterial, and Fungal Infections 515–524 (Elsevier, 2023).

2. Roy, S., Ghatak, D., Das, P. & BoseDasgupta, S. ESX secretion system: The gatekeepers of mycobacterial survivability and pathogenesis. Eur. J. Microbiol. Immunol. 10, 202–209 (2020).

3. Granados-Tristán, A. L. et al. ESX-3 secretion system in Mycobacterium: An overview. Biochimie (2023).

4. Mehra, A. et al. Mycobacterium tuberculosis type VII secreted effector EsxH targets host ESCRT to impair trafficking. PLoS Pathog. 9, e1003734 (2013).

5. Mittal, E. et al. Mycobacterium tuberculosis type VII secretion system effectors differentially impact the ESCRT endomembrane damage response. MBio 9, 10–1128 (2018).

6. Derrick, S. C. & Morris, S. L. The ESAT6 protein of Mycobacterium tuberculosis induces apoptosis of macrophages by activating caspase expression. Cell. Microbiol. 9, 1547–1555 (2007).

7. De Jonge, M. I. et al. ESAT-6 from Mycobacterium tuberculosis dissociates from its putative chaperone CFP-10 under acidic conditions and exhibits membrane-lysing activity. J. Bacteriol. 189, 6028–6034 (2007).

8. Boesen, H., Jensen, B. N., Wilcke, T. & Andersen, P. Human T-cell responses to secreted antigen fractions of Mycobacterium tuberculosis. Infect. Immun. 63, 1491–1497 (1995).

9. Demissie, A. et al. T-cell recognition of Mycobacterium tuberculosis culture filtrate fractions in tuberculosis patients and their household contacts. Infect. Immun. 67, 5967–5971 (1999).

10. Albrethsen, J. et al. Proteomic profiling of Mycobacterium tuberculosis identifies nutrient-starvation-responsive toxin–antitoxin systems. Mol. Cell. Proteomics 12, 1180–1191 (2013).

11. Ma, G. et al. Improving basic and membrane protein MS detection of the culture filtrate proteins from Mycobacterium tuberculosis H37Rv by biomimetic affinity prefractionation. Proteomics 17, 1600177 (2017).

12. Tucci, P., Portela, M., Chetto, C. R., González-Sapienza, G. & Marín, M. Integrative proteomic and glycoproteomic profiling of Mycobacterium tuberculosis culture filtrate. PloS One 15, e0221837 (2020).

13. Bell, C., Smith, G. T., Sweredoski, M. J. & Hess, S. Characterization of the Mycobacterium tuberculosis proteome by liquid chromatography mass spectrometry-based proteomics techniques: a comprehensive resource for tuberculosis research. J. Proteome Res. 11, 119–130 (2012).

14. Wang, Q. et al. PE/PPE proteins mediate nutrient transport across the outer membrane of Mycobacterium tuberculosis. Science 367, 1147–1151 (2020).

15. Leddy, O., White, F. M. & Bryson, B. D. Immunopeptidomics reveals determinants of Mycobacterium tuberculosis antigen presentation on MHC class I. Elife 12, e84070 (2023).

16. Lewinsohn, D. A. et al. Comprehensive definition of human immunodominant CD8 antigens in tuberculosis. Npj Vaccines 2, 8 (2017).

17. Ontiveros-Padilla, L. et al. CD4+ and CD8+ circulating memory T cells are crucial in the protection induced by vaccination with Salmonella Typhi porins. Microorganisms 9, 770 (2021).

18. Hermann, C., Karamchand, L., Blackburn, J. M. & Soares, N. C. Cell envelope proteomics of mycobacteria. J. Proteome Res. 20, 94–109 (2020).

19. Olson, M. G. et al. Proximity labeling to map host-pathogen interactions at the membrane of a bacterium-containing vacuole in Chlamydia trachomatis-infected human cells. Infect. Immun. 87, 10–1128 (2019).

20. Rodríguez-Ortega, M. J. “Shaving” live bacterial cells with proteases for proteomic analysis of surface proteins. in The Surfaceome 21–29 (Springer, 2018).

21. Mitra, A., Speer, A., Lin, K., Ehrt, S. & Niederweis, M. PPE surface proteins are required for heme utilization by Mycobacterium tuberculosis. MBio 8, 10–1128 (2017).

22. Ates, L. S. et al. Mutations in ppe38 block PE_PGRS secretion and increase virulence of Mycobacterium tuberculosis. Nat. Microbiol. 3, 181–188 (2018).

23. Dong, D. et al. PPE38 modulates the innate immune response and is required for Mycobacterium marinum virulence. Infect. Immun. 80, 43–54 (2012).

24. Becker, K. & Sander, P. Mycobacterium tuberculosis lipoproteins in virulence and immunity–fighting with a double-edged sword. FEBS Lett. 590, 3800–3819 (2016).

25. Su, H. et al. Mycobacterium tuberculosis PPE60 antigen drives Th1/Th17 responses via Toll-like receptor 2–dependent maturation of dendritic cells. J. Biol. Chem. 293, 10287–10302 (2018).

26. Jaisinghani, N. et al. Proteomics from compartment-specific APEX2 labeling in Mycobacterium tuberculosis reveals Type VII secretion substrates in the cell wall. Cell Chem. Biol. 31, 523–533 (2024).

27. Ahamed, M. et al. Optimized APEX2 peroxidase-mediated proximity labeling in fast-and slow-growing mycobacteria. in Methods in Enzymology vol. 664 267–289 (Elsevier, 2022).

28. Logsdon, M. M. & Aldridge, B. B. Stable regulation of cell cycle events in mycobacteria: Insights from inherently heterogeneous bacterial populations. Front. Microbiol. 9, 348921 (2018).

29. Götzke, H. et al. The ALFA-tag is a highly versatile tool for nanobody-based bioscience applications. Nat. Commun. 10, 4403 (2019).

30. Bosch, J. A., Chen, C. & Perrimon, N. Proximity-dependent labeling methods for proteomic profiling in living cells: An update. Wiley Interdiscip. Rev. Dev. Biol. 10, e392 (2021).

31. Xu, Y., Fan, X. & Hu, Y. In vivo interactome profiling by enzyme-catalyzed proximity labeling. Cell Biosci. 11, 27 (2021).

32. Sampson, S. L. Mycobacterial PE/PPE proteins at the host-pathogen interface. J. Immunol. Res. 2011, (2011).

33. Gong, Z. et al. Regulation of host cell pyroptosis and cytokines production by Mycobacterium tuberculosis effector PPE60 requires LUBAC mediated NF-κB signaling. Cell. Immunol. 335, 41–50 (2019).

34. Zhang, J. et al. PE12 interaction with TLR4 promotes intracellular survival of Mycobacterium tuberculosis by suppressing inflammatory response. Int. J. Biol. Macromol. 253, 127547 (2023).

35. Deng, W. et al. Mycobacterium tuberculosis PPE family protein Rv1808 manipulates cytokines profile via co-activation of MAPK and NF-κB signaling pathways. Cell. Physiol. Biochem. 33, 273–288 (2014).

36. Deng, W., Yang, W., Zeng, J., Abdalla, A. E. & Xie, J. Mycobacterium tuberculosis PPE32 promotes cytokines production and host cell apoptosis through caspase cascade accompanying with enhanced ER stress response. Oncotarget 7, 67347 (2016).

37. Meng, L. et al. PPE38 protein of Mycobacterium tuberculosis inhibits macrophage MHC class I expression and dampens CD8+ T cell responses. Front. Cell. Infect. Microbiol. 7, 68 (2017).

38. Song, H., Sandie, R., Wang, Y., Andrade-Navarro, M. A. & Niederweis, M. Identification of outer membrane proteins of Mycobacterium tuberculosis. Tuberculosis 88, 526–544 (2008).

39. Målen, H., Pathak, S., Søfteland, T., De Souza, G. A. & Wiker, H. G. Definition of novel cell envelope associated proteins in Triton X-114 extracts of Mycobacterium tuberculosis H37Rv. BMC Microbiol. 10, 1–11 (2010).

40. De Souza, G. A., Leversen, N. A., Målen, H. & Wiker, H. G. Bacterial proteins with cleaved or uncleaved signal peptides of the general secretory pathway. J. Proteomics 75, 502–510 (2011).

41. Sharma, T. et al. The Mycobacterium tuberculosis PE_PGRS protein family acts as an immunological decoy to subvert host immune response. Int. J. Mol. Sci. 23, 525 (2022).

42. Nair, S. et al. The PPE18 of Mycobacterium tuberculosis interacts with TLR2 and activates IL-10 induction in macrophage. J. Immunol. 183, 6269–6281 (2009).

43. Skeiky, Y. A. et al. Differential immune responses and protective efficacy induced by components of a tuberculosis polyprotein vaccine, Mtb72F, delivered as naked DNA or recombinant protein. J. Immunol. 172, 7618–7628 (2004).

44. Dillon, D. C. et al. Molecular characterization and human T-cell responses to a member of a novel Mycobacterium tuberculosis mtb39 gene family. Infect. Immun. 67, 2941–2950 (1999).

45. Tait, D. R. et al. Final analysis of a trial of M72/AS01E vaccine to prevent tuberculosis. N. Engl. J. Med. 381, 2429–2439 (2019).

46. Van Der Meeren, O. et al. Phase 2b controlled trial of M72/AS01E vaccine to prevent tuberculosis. N. Engl. J. Med. 379, 1621–1634 (2018).

47. De Valliere, S., Abate, G., Blazevic, A., Heuertz, R. & Hoft, D. Enhancement of innate and cell-mediated immunity by antimycobacterial antibodies. Infect. Immun. 73, 6711–6720 (2005).

48. Chen, T. et al. Association of human antibodies to arabinomannan with enhanced mycobacterial opsonophagocytosis and intracellular growth reduction. J. Infect. Dis. 214, 300–310 (2016).

49. McIntyre, S., Warner, J., Rush, C. & Vanderven, H. A. Antibodies as clinical tools for tuberculosis. Front. Immunol. 14, 1278947 (2023).

50. Li, H. & Javid, B. Antibodies and tuberculosis: finally coming of age? Nat. Rev. Immunol. 18, 591–596 (2018).

51. Correia-Neves, M., Sundling, C., Cooper, A. & Källenius, G. Lipoarabinomannan in active and passive protection against tuberculosis. Front. Immunol. 10, 477025 (2019).

52. Watson, A. et al. Human antibodies targeting a Mycobacterium transporter protein mediate protection against tuberculosis. Nat. Commun. 12, 602 (2021).

53. Spratt, J. M., Ryan, A. A., Britton, W. J. & Triccas, J. A. Epitope-tagging vectors for the expression and detection of recombinant proteins in mycobacteria. Plasmid 53, 269–273 (2005).

54. Guo, J. et al. The development of proximity labeling technology and its applications in mammals, plants, and microorganisms. Cell Commun. Signal. 21, 269 (2023).

55. Steinhäuser, C. et al. Lipid-Labeling facilitates a novel magnetic isolation procedure to characterize pathogen-containing phagosomes. Traffic 14, 321–336 (2013).

56. Hoffmann, E., Machelart, A., Song, O.-R. & Brodin, P. Proteomics of mycobacterium infection: moving towards a better understanding of pathogen-driven immunomodulation. Front. Immunol. 9, 86 (2018).

57. Ferrari, G., Langen, H., Naito, M. & Pieters, J. A coat protein on phagosomes involved in the intracellular survival of mycobacteria. Cell 97, 435–447 (1999).

58. Clemens, D. L., Lee, B.-Y. & Horwitz, M. A. Deviant expression of Rab5 on phagosomes containing the intracellular pathogens Mycobacterium tuberculosis and Legionella pneumophila is associated with altered phagosomal fate. Infect. Immun. 68, 2671–2684 (2000).

59. Kasmapour, B., Gronow, A., Bleck, C. K., Hong, W. & Gutierrez, M. G. Size-dependent mechanism of cargo sorting during lysosome-phagosome fusion is controlled by Rab34. Proc. Natl. Acad. Sci. 109, 20485–20490 (2012).

60. Chandra, P. et al. Mycobacterium tuberculosis inhibits RAB7 recruitment to selectively modulate autophagy flux in macrophages. Sci. Rep. 5, 1–10 (2015).

61. Reddington, S. C. & Howarth, M. Secrets of a covalent interaction for biomaterials and biotechnology: SpyTag and SpyCatcher. Curr. Opin. Chem. Biol. 29, 94–99 (2015).

