## Supplementary figures and images for "Protease shaving of *Mycobacterium tuberculosis* facilitates vaccine antigen discovery and delivery of novel cargoes to the Mtb surface"

### Supplemental Figure 1

#### Supplemental Figure 1

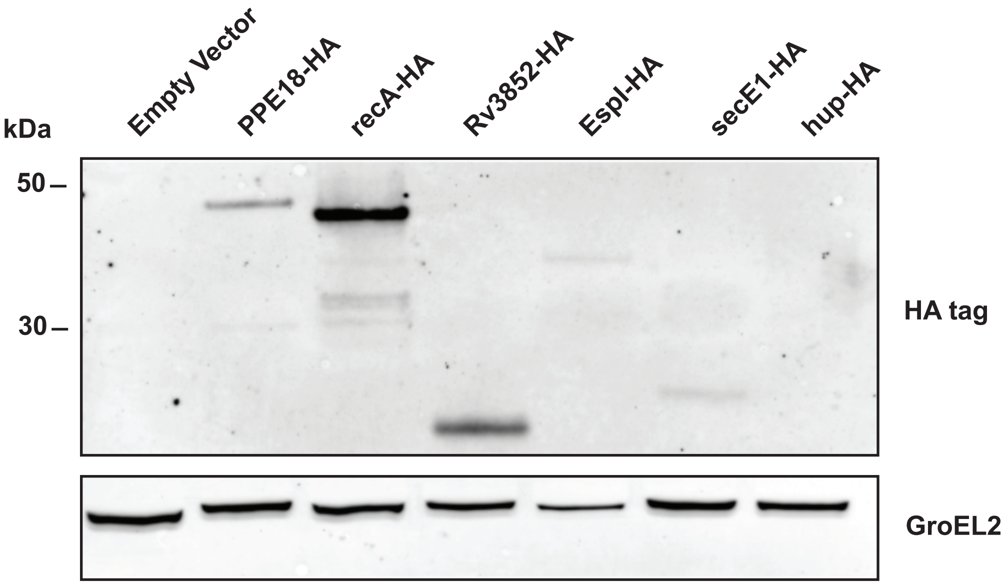

### Supplemental Figure 2

#### Supplemental Figure 2

(A)

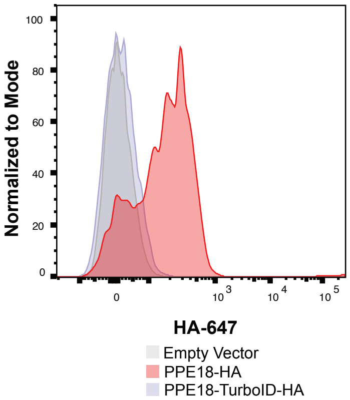


(B)

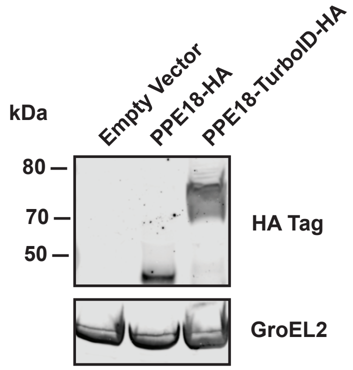

### Supplemental Figure 3

##### Supplemental Figure 3

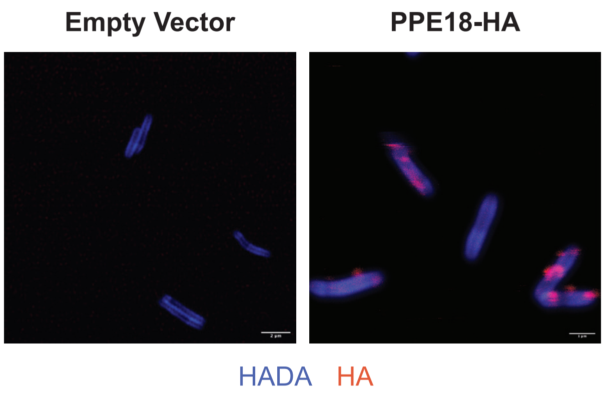

### Supplemental Figure 4

**Supplemental Figure 4**

**(A)**

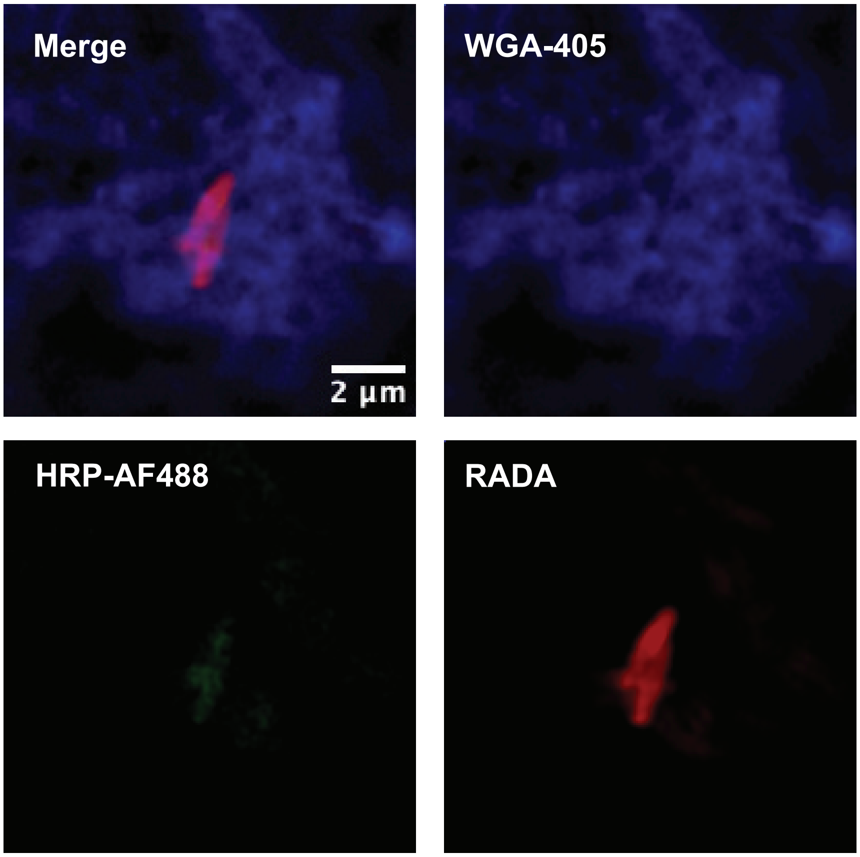


**(B)**

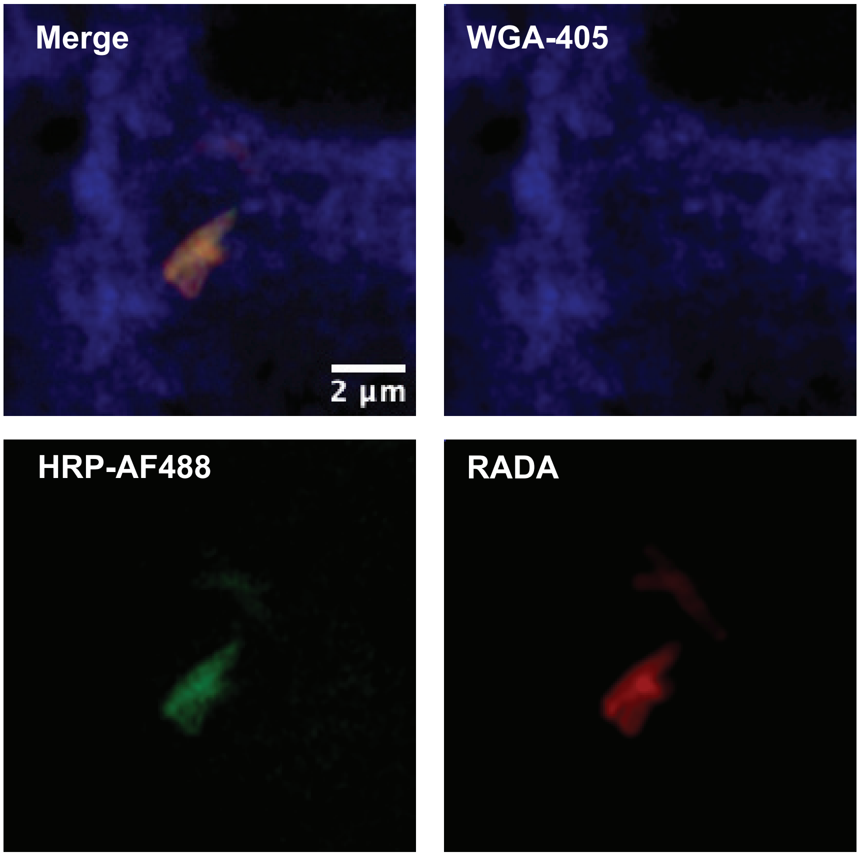
